# Baseline Semantic Fluency Is Associated with Six-Year Progression to Mild Cognitive Impairment in Middle-Aged Men

**DOI:** 10.1101/659417

**Authors:** Daniel E. Gustavson, Jeremy A. Elman, Matthew S. Panizzon, Carol E. Franz, Jordan Zuber, Mark Sanderson-Cimino, Chandra A. Reynolds, Kristen C. Jacobson, Hong Xian, Amy J. Jak, Rosemary Toomey, Michael J. Lyons, William S. Kremen

**Affiliations:** Department of Otolaryngology, Vanderbilt University Medical Center, Nashville, TN. Address: 1215 21^st^ Ave South (10420B MCE), Nashville, TN 37232.; Department of Psychiatry, Center for Behavior Genetics of Aging, University of California, San Diego, La Jolla, CA; Department of Psychology, San Diego State University, San Diego, CA; Department of Psychology, University of California, Riverside, Riverside, CA; Department of Psychiatry and Behavioral Neuroscience, University of Chicago, Chicago, IL; Department of Epidemiology and Biostatistics, St. Louis University and Clinical Epidemiology Center, Veterans Affairs St. Louis Healthcare System, St. Louis, MO; Department of Psychiatry, University of California, San Diego, La Jolla, CA and Psychology Service and Center of Excellence for Stress and Mental Health, Veterans Affairs San Diego Healthcare System, La Jolla, CA; Department of Psychological and Brain Sciences, Boston University, Boston, MA; Department of Psychiatry, Center for Behavior Genetics of Aging, University of California, San Diego, La Jolla, CA and Center of Excellence for Stress and Mental Health, Veterans Affairs San Diego Healthcare System, La Jolla, CA

**Author notes:** The corresponding author, Daniel E. Gustavson, conducted all statistical analyses.

**Keywords:** MCI, Alzheimer’s disease, cognitive aging, memory, executive function

## Abstract

**Objective:** Test the hypothesis that individual differences in episodic memory and verbal fluency in cognitively normal middle-aged adults will predict progression to amnestic MCI after 6 years.

**Method:** The analysis sample included 842 male twins who were cognitively normal at baseline (M=56 years), completed measures of episodic memory and verbal fluency at baseline and again 6 years later (M=62 years).

**Results:** Poor episodic memory predicted progression to both amnestic MCI (OR=4.42, 95% CI [2.44, 10.60]) and non-amnestic MCI (OR=1.92, 95% CI [1.32, 3.44]). Poor semantic verbal fluency also independently predicted progression to amnestic MCI (OR=1.86, 95% CI [1.12, 3.52]). In the full sample, a semantic-specific fluency latent variable at wave 1 (which controls for letter fluency) predicted change in episodic memory at wave 2 (*β*=.13), but not vice-versa (*β*=.04). Associations between episodic memory and verbal fluency factors were primarily explained by genetic, rather than environmental, correlations.

**Conclusions:** Among individuals who were cognitively normal at wave 1, episodic memory moderately-to-strongly predicted progression to MCI at average age 62, emphasizing the fact that there is still meaningful variability even among cognitively normal individuals. Episodic memory, which is typically a primary focus for AD risk, declined earlier and more quickly than fluency. However, semantic fluency at average age 56 predicted 6-year change in memory as well as progression to amnestic MCI even after accounting for baseline memory performance. These findings emphasize the utility of memory and fluency measures in early identification of AD risk.

## Introduction

Although the amyloid/tau/neurodegeneration (A/T/[N]) framework is agnostic regarding the sequence of biomarker progression, it is generally assumed that biomarker positivity is present before cognitive manifestations^1^. Amyloid beta and tau biomarkers indicate increased risk because they reflect underlying disease pathophysiology, but it does not necessarily follow that–with currently available detection techniques–these biomarkers will be the earliest predictors of progression to AD. In non-demented older adults, including those who already have mild cognitive impairment (MCI), cognitive markers–particularly episodic memory–do as well or better than AD biomarkers^2-5^. Semantic fluency impairment has also been associated with MCI and progression to AD in older adults^6-12^. However, because the AD process begins decades before dementia onset, researchers have recommended greater emphasis on earlier identification of risk^13,14^.

With that in mind, there is still substantial variability in cognitive function even in cognitively normal, middle-aged adults. To address the need for early identification, we examined the ability of cognitive measures to predict progression to MCI in a 6-year follow-up of cognitively normal, community-dwelling men in their 50s who were rigorously defined as cognitively normal at baseline. We addressed the following questions: 1) Does episodic memory predict progression to MCI? 2) Does semantic fluency predict progression to MCI after accounting for episodic memory? 3) Does episodic memory predict change in semantic fluency or vice versa? 4) To what extent do shared genetic influences underlie associations between semantic fluency and episodic memory?

## Method

### Participants

The primary analyses involving MCI progression (questions 1 and 2) were based on 842 individuals from the longitudinal Vietnam Era Twin Study of Aging (VETSA) project who were cognitively normal at wave 1 and returned to complete the wave 2 assessment approximately six years later. All participants were recruited randomly from the Vietnam Era Twin Registry from a previous study^15^. All served in the United States military at some time between 1965 and 1975; nearly 80% did not serve in combat or in Vietnam^16,17^. Participants are generally representative of American men in their age group with respect to health and lifestyle characteristics^18^. See Table 1 for demographic characteristics of the subsample of 842 individuals (and Table S1 for the full sample), which includes the descriptive statistics for all covariates in the analyses involving MCI. All participants provided informed consent, and all procedures were approved the Institutional Review Board of participating institutions.

**Table 1.** Demographic Characteristics of the Sample and Covariates Included in Analyses Involving Mild Cognitive Impairment (MCI)

| Demographic Variable | M | SD | Range | *p*<br>(CN vs.<br>aMCI) | *p*<br>(CN vs.<br>nMCI) | Overall *p*<br>(CN vs.<br>aMCI vs.<br>nMCI) |
| --- | --- | --- | --- | --- | --- | --- |
| All Subjects |  |  |  |  |  |  |
| Lifetime Education | 14.0 | 2.13 | 8, 20 | 0.229 | 0.168 | 0.210 |
| Ethnicity (% white non-Hispanic) | 91.0 | - | - | 0.793 | 0.743 | 0.325 |
| ApoE status (% e4 positive) | 30.5 | - | - | 0.499 | 0.093 | 0.203 |
| Wave 1 |  |  |  |  |  |  |
| Age | 55.8 | 2.46 | 51.08, 60.67 | 0.136 | 0.008 | 0.004 |
| Depression Symptoms | 7.17 | 7.17 | 0, 45 | 0.809 | 0.818 | 0.955 |
| Diabetes (% yes) | 10.3 | - | - | 0.747 | 0.420 | 0.554 |
| Hypertension (% yes) | 59.4 | - | - | 0.259 | 0.140 | 0.248 |
| Wave 2 |  |  |  |  |  |  |
| Age | 61.5 | 2.41 | 56.50, 66.50 | 0.098 | 0.012 | 0.005 |
| Age Interval (wave 2 - wave 1) | 5.8 | 0.69 | 4.25, 9.42 | 0.607 | 0.510 | 0.682 |

*Note:* N = 842 individuals who were cognitively normal at wave 1 and returned for wave 2 assessments. Column 6 and 7 display the *p* value for comparisons between the individuals who remained cognitively normal (N=762) and either progressed to amnesic MCI (aMCI; N=42; Column 6) or nonamnesic MCI (nMCI; N=38; Column 7) at wave 2. The final column displays the 2 degree of freedom comparison between all three groups. Lifetime education was the number of years of school completed. Depression symptoms were measured with the Center for Epidemiologic Studies – Depression Scale<sup>36</sup>, with scores above 15 indicating risk for clinical depression. Descriptive statistics for the full sample are displayed in Table S1.

There were 1237 individuals who completed the protocol at wave 1. Of these, 107 (8.6%) were excluded from the analysis of progression to MCI because they had MCI at wave 1. This left 1130 (91.4%) CN individuals, 906 (80.2%) of whom returned for wave 2. Of those 906, 64 were excluded for the following reasons: missing covariates (57); missing fluency data (1); MCI with language domain impairment (which includes fluency) at wave 2 (6). This left 842 (92.9% of 906 and 74.5% of the total CN individuals at wave 1) for the analyses of progression to MCI. See supplemental Figure S1 for a flowchart describing this sample breakdown.

Additional analyses involving the full sample (questions 3 and 4) made use of data from all VETSA participants: 1484 total individuals. At wave 2, participants included 1257 individuals: 1013 returnees, 191 attrition replacement subjects who were in the age range of returnees, and 53 attrition replacements who were in the age range of participants at wave 1. The final 53 attrition replacements were analyzed with the wave 1 subjects. Despite some missing data, all available subjects were included because structural equation models (i.e., phenotypic cross-lagged and genetic twin models) use all available data. This provides more accurate and less biased estimates of variance, thus making for more accurate factor loadings and estimates of genetic/environmental influences at each age. Thus, the final sample included 1290 individuals at mean age 56 (367 monozygotic [MZ] twin pairs, 273 dizygotic [DZ] twin pairs, and 10 unpaired twins), and 1204 individuals at mean age 62 (332 MZ twin pairs, 234 DZ twin pairs, and 72 unpaired twins).

### Measures

Development of latent variable measures of episodic memory, verbal fluency, and other cognitive variables, and MCI diagnoses has been reported in our earlier work^19-22^. These measures are summarized briefly here. Additionally, all cognitive measures at the second wave of testing were adjusted to account for the fact that many of the subjects had encountered the tasks before^20^. Although practice effects were small and often non-significant, it is important to correct for them as ignoring these small differences result in the underdiagnoses of MCI in this sample^20^.

### Episodic memory

Memory was measured with the Logical Memory (LM) and Visual Reproductions (VR) subtests of the Wechsler Memory Scale-III^23^, and the California Verbal Learning Test-II (CVLT)^24^. The dependent measures were the LM and VR delayed recall scores, and the CVLT long delay free recall score. These measures were combined to create a latent factor.

### Verbal fluency

Fluency was measured with 6 verbal fluency subtests from the Delis-Kaplan Executive Function System (D-KEFS)^25^. Subjects performed the letter subtests (*F, A*, and *S*), followed by semantic subtests (*Animals, Boys’ Names*, and *Fruits/Furniture*). Dependent measures were the number of words generated in 60 seconds. We utilized a latent variable model of verbal fluency that highlights unique variance in semantic fluency^26^. This model decomposes the variance in the 6 fluency subtests into two latent factors: a <u>general fluency</u> factor captures variance across letter and semantic subtests, and a <u>semantic-specific</u> factor captures the remaining variance shared among semantic subtests not captured by the general factor. To be comparable with the other measures, we used the number of words on Fruits/furniture switching condition, ignoring switching ability. This was validated in the previous work^26^, which also showed that removal of this subtest has little impact on the factor structure of verbal fluency. Performance on both factors was explained mostly by genetic influences at wave 1 and wave 2 and demonstrated strong phenotypic and genetic correlations over this six-year interval^26^.

### Mild cognitive impairment

MCI was diagnosed using the Jak-Bondi actuarial/neuropsychological approach^27-29^. Impairment in a cognitive domain was defined as having at least two tests >1.5 SDs below the age- and education-adjusted normative means after accounting for “premorbid” cognitive ability by adjusting neuropsychological scores for performance on a test of general cognitive ability that was taken at a mean age of 20 years. The adjustment for age 20 cognitive ability ensures that the MCI diagnosis is capturing a decline in function rather than long-standing low ability. The validity of the VETSA MCI diagnoses is supported by previous studies^27-29^ and in the present sample by evidence of reduced hippocampal volume in those diagnosed with amnestic MCI^30^. Higher AD polygenic risk scores were associated with significantly increased odds of MCI^19^, indicating that our diagnosis is genetically-related to AD.

Because we were interested in transition to MCI, analyses involving incident MCI at wave 2 only included individuals who were cognitively normal at wave 1 and had data for all covariates (see supplement Figure S1). Of the 842 returnees meeting this criterion, 42 (5.0%) progressed to amnestic MCI, and 38 (4.5%) progressed to non-amnestic MCI. An additional 6 individuals who had MCI with language domain impairment (3 of which were also amnestic), were assigned missing values for all analyses involving MCI. This was done to prevent biasing estimates of the association between verbal fluency and MCI because letter and semantic fluency comprised the language domain. However, odds ratios were nearly identical if these individuals were all included.

### Other cognitive measures (wave 1 only)

To examine how other cognitive abilities account for the overlap between episodic memory and semantic fluency, a latent factor for vocabulary was created using the vocabulary subtest of the Wechsler Abbreviated Scale of Intelligence (WASI)^31^ and the multiple-choice vocabulary subtest of the Armed Forced Qualification Test^32,33^. Two executive function latent factors were included based on our previous work^34^. The first factor (Common Executive Function) captures common variance across 6 tasks assessing inhibition, shifting, and working memory^21^. The second latent factor (Working Memory-Specific) captures covariance among working memory tasks not already captured by the common factor.

### Data Analysis

Phenotypic analyses involving MCI were conducted with mixed effects logistic regression using the lme4 package in R version 3.3.1^35^. The lme4 package uses list-wise deletion with missing observations, and reports profile-based 95% confidence intervals (95% CIs). In all models, pair ID was included as a random effect to account for the clustering of data within families. We also controlled for wave 1 age, diabetes, hypertension, depression symptoms (based on Center for Epidemiologic Studies–Depression scale)^36^, the time interval between wave 1 and wave 2, *ApoE* status (ε4+ alleles vs. ε4-), and years of education. Post-hoc analyses indicated no evidence that prediction of later MCI differed in MZ versus DZ twins (i.e., no memory x zygosity or fluency x zygosity interactions), so these interaction terms were excluded in all models presented. Additional post hoc analyses revealed that the pattern of results were nearly identical without controlling for any covariates (see supplement Table S2).

Additional phenotypic regression and cross-lagged analyses in the full sample were conducted using Mplus Version 7^37^. Genetic analyses were conducted using the structural equation modeling package OpenMx in R^38^. Both programs account for missing observations using a full-information maximum likelihood approach. Model fit for these structural equation models was determined using *χ*^2^ tests, the Root Mean Square Error of Approximation (RMSEA), and the Comparative Fit Index (CFI). Good fitting models had *χ*^2^ values less than 2 times the degrees of freedom, RMSEA values<.06, and CFI values>.95^39^. Additionally, good fitting genetic models did not fit significantly worse than a full Cholesky decomposition of all measures, a common baseline model in twin studies. Significance of individual parameters was established with *χ*^2^ difference tests and standard-error based (Mplus) or likelihood-based (OpenMx) 95% CI.

Genetically-informed models were based on the standard assumptions in twin designs, which decompose variance in phenotype (and covariance among phenotypes) into three sets of influences: additive genetic influences (A), shared environmental influences (C), and nonshared environmental influences (E). Additive genetic influences are correlated at 1.0 in MZ twin pairs and 0.5 in DZ twin pairs because MZ twins share all their alleles identical-by-descent and DZ twins share, on average, 50% of their alleles identical-by-descent. Shared environmental influences (C), which are environmental influences that make twins in a pair more similar, are correlated at 1.0 in both MZ and DZ twins. Nonshared environmental influences (E), which are environmental influences that make twins in a pair dissimilar, are correlated at 0.0 in both MZ and DZ twin pairs, by definition. We also assume equal means and variances within pairs and across zygosity. These assumptions for univariate analyses apply to multivariate cases and to situations where phenotypic correlations between constructs are decomposed into their genetic (*r*_genetic_) and nonshared environmental components (*r*_environmental_). Shared environmental influences on verbal fluency factors, and their correlations with episodic memory, were not estimated based on our previous work showing no evidence for shared environment on verbal fluency^26^.

## Data Availability

VETSA data are publicly available, with restrictions. Information regarding data access can be found at our website: https://medschool.ucsd.edu/som/psychiatry/research/VETSA/Pages/default.aspx

## Results

### Descriptive Statistics

Demographic and clinical characteristics are displayed in Table 1. There were no significant differences between cognitively normal and MCI groups in any of these characteristics, except that individuals who progressed to amnestic MCI were older than those who remained cognitively normal at both wave 1 (*p*=.008) and wave 2 (*p*=.012). Descriptive statistics for fluency and memory measures are displayed in the supplement (Table S3).

### Predicting Progression to MCI

Table 2 displays the longitudinal logistic regression analyses. Progression to amnestic MCI (Table 2A) or non-amnestic MCI (Table 2B) at wave 2 is predicted by general fluency, semantic-specific and episodic memory factor scores at wave 1, as well as the covariates. Higher odds ratios indicate increased odds of progression to MCI (estimated for −1 *SD* decrease on the original factor score scale). Supplement Table S4 displays similar analyses of progression to any MCI (i.e., amnestic and nonamnestic MCI collapsed into a single group).

**Table 2.** Logistic Regression for Mild Cognitive Impairment (MCI) at Wave 2 Predicted by Fluency and Memory Factor Scores at Wave 1

| Dependent Variable | A) No MCI at Wave 2<br>(N=762) vs. Amnestic MCI<br>at Wave 2 (N=42) |  | B) No MCI at Wave 2<br>(N=762) vs. Non-Amnestic<br>MCI at Wave 2 (N=38) |  |
| --- | --- | --- | --- | --- |
|  | OR | 95% CI | OR | 95% CI |
| *Cognitive Factor Scores (Wave 1)* |  |  |  |  |
| General Fluency | 1.29 | [0.79, 2.35] | 1.29 | [0.82, 2.12] |
| Semantic-Specific | **1.86** | **[1.12, 3.52]** | 0.92 | [0.59, 1.42] |
| Episodic Memory | **4.42** | **[2.44, 10.60]** | **1.92** | **[1.32, 3.34]** |
| *Covariates* |  |  |  |  |
| Age (wave 1) | 1.28 | [0.76, 2.32] | **1.77** | **[1.13, 3.05]** |
| Age Interval (wave 2 - wave 1) | 1.41 | [0.87, 2.44] | 0.99 | [0.58, 1.57] |
| Depression (wave 1) | 0.92 | [0.56, 1.47] | 1.01 | [0.66, 1.52] |
| ApoE ε4+ | 1.49 | [0.54, 4.95] | 0.47 | [0.13, 1.25] |
| Diabetes (wave 1) | 1.05 | [0.21, 4.47] | 1.55 | [0.43, 4.94] |
| Hypertension (wave 1) | 1.35 | [0.52, 4.28] | 1.69 | [0.72, 4.48] |
| Years of Education | 0.96 | [0.57, 1.65] | 0.82 | [0.49, 1.29] |

*Note:* Significant odds ratios (ORs) are displayed in bold ( $p < .05$ ). Factor scores for verbal fluency and episodic memory were scored and standardized so the odds ratio indicate the increase in odds of converting to amnestic MCI (A) or non-amnestic MCI (B) at -1 *SD* for that variable. Individuals with non-amnestic MCI were excluded from analyses of amnestic MCI (A) and vice-versa. Measures of age, depression, and years of education were also standardized, but not reverse scored. Odds ratios for ApoE status, diabetes, and hypertension reflect increase in odds for having an ε4 allele, diabetes, or hypertension, respectively. Both analyses also included a random effect of family (Table 2A: *SD* = 1.91, Table 2B: *SD* = 1.52). CI = Confidence interval.

Results indicated that episodic memory at wave 1 strongly predicted progression to amnestic MCI at wave 2, even when controlling for both verbal fluency factors, OR=4.42, *p*<.001, 95% CI [2.44, 10.60]. Poor semantic-specific fluency was also associated with nearly double the odds of being diagnosed with amnestic MCI at wave 2 even when controlling for episodic memory, OR=1.86, *p*=.025, 95% CI [1.12, 3.52]. Poor general fluency was not associated with increased odds of amnestic MCI, OR=1.29, *p*=.332, 95% CI [0.79, 2.35]. Interestingly, poor episodic memory was also associated with progression to non-amnestic MCI, OR=1.92, *p*=.009, 95% CI [1.21, 3.34]. Neither fluency factor predicted progression to non-amnestic MCI, both *p*s>.274.

### Longitudinal Associations between Episodic Memory and Verbal Fluency

We next fit a phenotypic cross-lagged model using fluency and memory data from both waves in the full sample. As seen in Figure 1, memory at mean age 62 was significantly predicted by semantic-specific fluency at mean age 56, *β*=.13, *p*=.035, 95% CI [.01, .25], even after accounting for memory at age 56, *β*=.85, *p*<001, 95% CI [.74, .98]. In contrast, memory at age 56 did not predict semantic-specific fluency at age 62, *β*=.04, *p*=.582, 95% CI [-.09, .16]. The general fluency factor also did not predict later memory (or vice versa).

**Figure 1:**
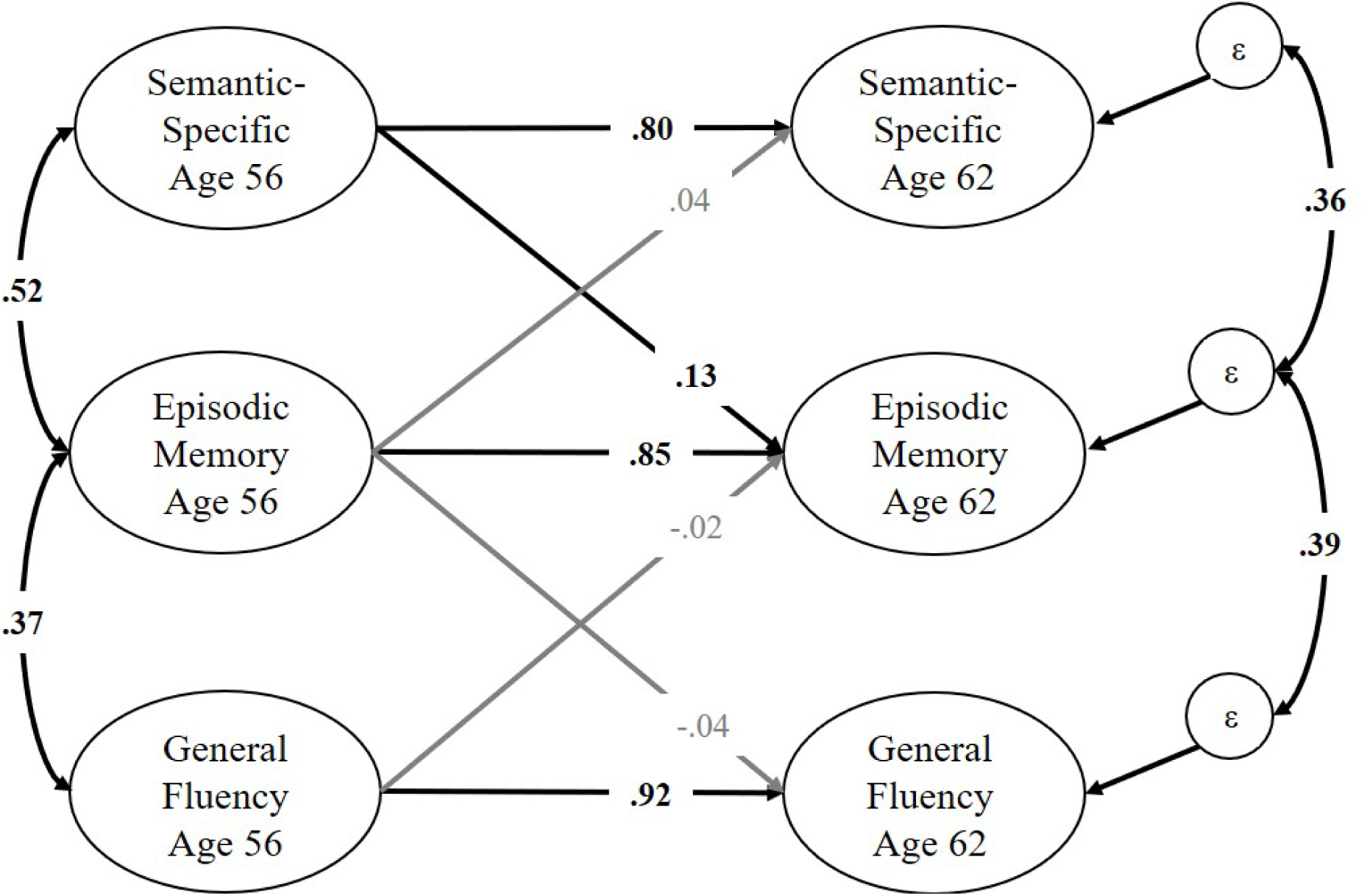
Phenotypic cross-lagged model of verbal fluency factors and episodic memory. Both fluency latent factors are correlated with episodic memory at wave 1 (age 56) and predict episodic memory at wave 2 (age 62). Episodic memory at wave 1 also predicts the fluency factors at wave 2. Each factor at wave 1 also predicts itself at wave 2, and there are residual correlations between the residual variances (ε) on the fluency factors at wave 2 and the episodic memory factor at wave 2. Not pictured are the factor loadings on individual tasks (which were equated across time and similar in magnitude to those displayed in Figure 2) or residual correlations between all observed variables across time (e.g., *Animals* at wave 1 correlated with *Animals* at wave 2). Significant paths and correlations are displayed in bold, with black text and lines (*p* < .05). The semantic-specific factor is the only factor to significantly predict another construct at the second wave (*β* = .13).

### Biometric Models of Cognitive Abilities

In a biometric twin models of the age 56 and 62 data (Figure 2), the phenotypic correlations between memory and semantic-specific fluency were explained primarily by genetic influences, *r*_genetic_=.65, 95% CI [.46, .88] at age 56, *r*_genetic_=.73, 95% CI [.58, .94] at age 62, explaining 87% (age 56) and 86% (age 62) of the total phenotypic correlations. There was also a significant nonshared environmental correlation between episodic memory and semantic-specific fluency at age 62 only, *r*_environmental_=.28, 95% CI [.01, .55]. This environmental correlation was nonsignificant at age 56, but similar in magnitude, *r*_environmental_=.25, 95% CI [-.08, .60]. Genetic correlations explained 92% (age 56, *r*_genetic_=.50, 95% CI [.36, .70]) and 94% (age 62, *r*_genetic_=.36, 95% CI [.25, .50]) of the correlation between memory and general fluency.

**Figure 2:**
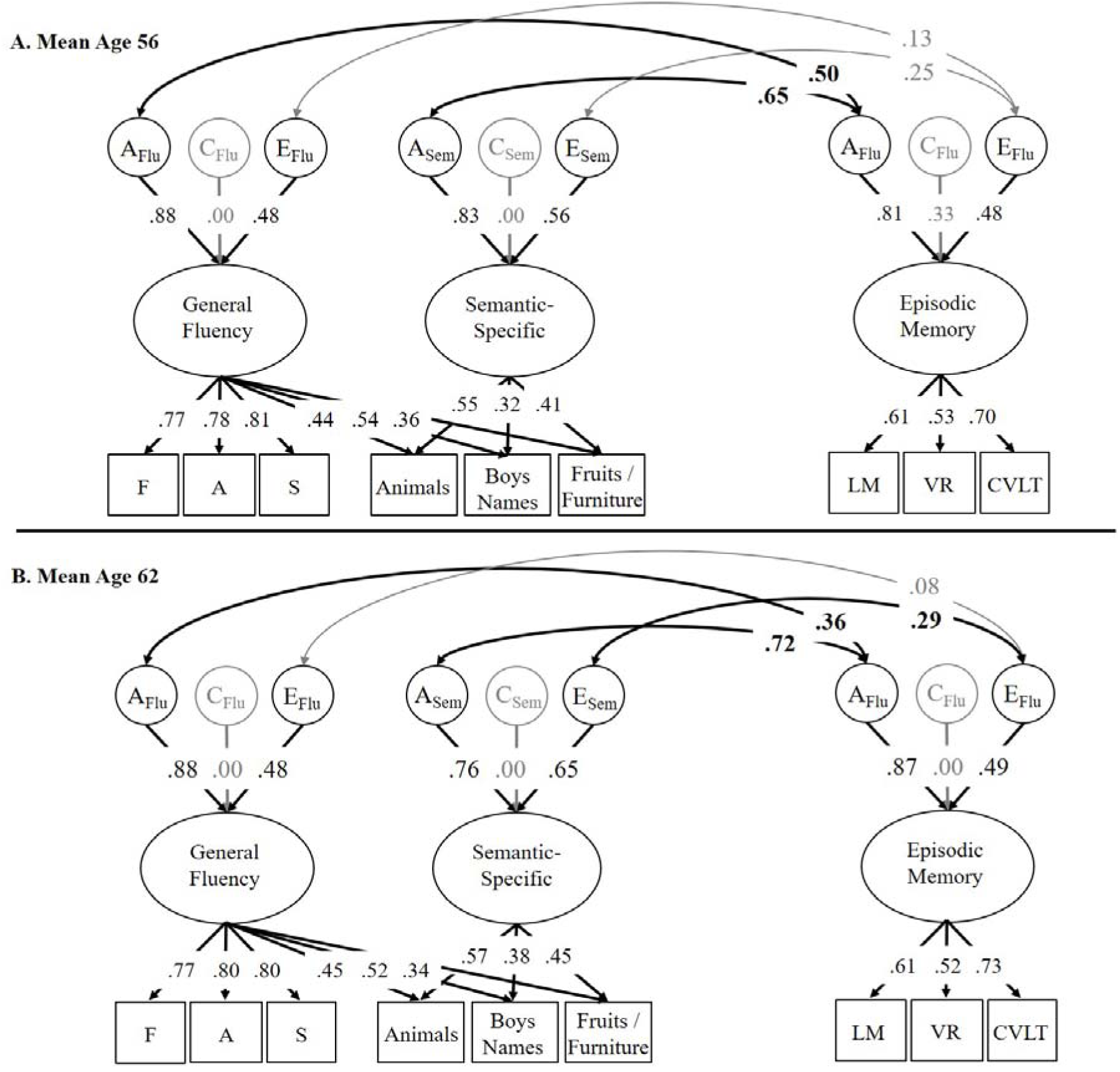
Full correlational models of the genetic (A), shared environmental (C), and nonshared environmental (E) influences on the general fluency, semantic-specific, and episodic memory latent variables at wave 1 (mean age 56; top) and wave 2 (mean age 62; bottom). Ellipses indicate latent variables and rectangles indicate measured variables. Significant factor loadings are displayed in bold, with black text and lines (*p* < .05). Variation explained by latent factors can be computed by squaring the factor loadings.

Using only the age 56 data, we fit a regression model (supplemental Figure S2) in which the 2 fluency factors were regressed on factors for memory, vocabulary, and the 2 executive function factors. Although semantic-specific fluency was positively correlated with episodic memory (*r*=.46, *p*<.001), vocabulary (*r*=.19, *p*<.001), and common executive function (*r*=.20, *p*=.003), only the association with memory remained significant in the regression model, *β*=.60, *p*<.001. Conversely, the general fluency factor was correlated with all other cognitive abilities (*r*s=.32 to .50) including memory, *r*=.38, *p*<.001, but this association with memory was nonsignificant in the regression model, *β*=.04, *p*=.538. These findings suggest that the genetic correlations between memory and general fluency were likely explained by their overlap with other cognitive abilities, but the association between episodic memory and semantic-specific fluency reflects unique genetic associations between these constructs.

## Discussion

These findings highlight the importance of episodic memory and semantic fluency as risk factors for cognitive decline and MCI. Among middle-aged men (ages 51-60) who were cognitively normal at wave 1, both measures predicted amnestic MCI 6 years later. Importantly, our previous results showing that higher AD polygenic risk scores were associated with significantly increased odds of having MCI in this sample provide support for this diagnosis being AD-related MCI^19^. The current study demonstrated that baseline semantic-specific fluency predicted progression to amnestic MCI even after controlling for baseline memory. In other previous work, we found that there was no change in the semantic-specific factor (*d*=-.01) over this 6-year interval^26^, yet in the current study it did independently predict episodic memory 6 years later. Thus, semantic-specific fluency is associated with episodic memory, but it does not necessarily change at the same time or rate as episodic memory. Nevertheless, it does add to the prediction of memory and progression to amnestic MCI beyond the predictive value of baseline memory itself.

With respect to amnestic MCI, the focus is naturally on episodic memory, but our results indicate that relatively poor semantic-specific fluency performance among cognitively normal adults may precede decline in episodic memory. The associations between semantic-specific fluency and episodic memory were driven by genetic influences at both waves. Furthermore, these genetic influences may overlap with those for AD. Additional research examining the relationship of cognitive and biomarker trajectories may determine the mechanisms driving this ordering of decline. For example, the functional networks contributing to these cognitive processes may be affected at different points in the aging or disease process due to the topographical spread of pathology or the vulnerability of particular brain regions to pathology. Studies utilizing molecular imaging of AD pathology (i.e., tau and amyloid PET) will be needed to further clarify the nature of these relationships. Regardless, these findings demonstrate that semantic fluency tests can be useful indicators of later memory impairment.

Although the prediction was not as strong as that for amnestic MCI, episodic memory at wave 1 also predicted non-amnestic MCI at wave 2, suggesting it can also be a predictor of cognitive decline in other domains (except fluency). Some of this prediction may also be due to AD pathology, where both memory and other cognitive abilities are declining together (but with memory not yet reaching criteria for amnestic MCI). However, this association may simply reflect other non-pathological changes in aging. Specifically, memory is more strongly correlated with general cognitive ability than verbal fluency in this sample (especially the semantic-specific factor, which adjusts for letter fluency)^40,41^, and general cognitive ability may be driving age-related changes across all domains^42,43^. Studies with multiple longitudinal assessments, especially those that can elucidate the relationship between biomarker and cognitive trajectories, will be useful in clarifying the causal mechanisms domain-specific changes.

Although biomarkers are necessary for biologically-based diagnosis, and are important for identifying individuals at greatest risk for cognitive decline and dementia, several studies have shown that neuropsychological tests are often better and earlier predictors of progression to AD than biomarkers^3-5,44-46^. One reason may be that current techniques for measuring biomarkers are not necessarily able to detect the earliest stages of disease progression. For example, beta-amyloid accumulates slowly for as much as 2 decades before symptom onset, but PET ligands have high affinity only for later stage neuritic beta-amyloid plaques^47,48^. Continuing the search for earlier biomarker detection is critically important. Meanwhile, the present results suggest that measures of memory and fluency are effective early indicators of risk for progression to amnestic MCI given that participants were only in their 50s at the baseline assessment. They can also be completed in a short amount of time at little expense. Fluency and memory measures may be combined with biomarkers to further improve prediction of MCI and AD. Biomarker data will improve determination of specificity for AD-related deficits, and fluency and memory measures may also be useful as screening tools for identifying individuals who should be followed up with biomarker testing.

### Strengths and Limitations

This sample comprised primarily white, non-Hispanic men, so these findings may not generalize across sex and race/ethnicity. It will also be important for future work to directly compare these odds ratios with those for biomarkers (preferably in the same model) and to examine whether neuropsychological tests more strongly predict progression to MCI in subsets of individuals (e.g., who are already amyloid positive). Finally, in most longitudinal studies, attriters tend to have lower cognitive ability than the returnees. In the present study, dropouts had significantly lower memory factor scores at wave 1, *p*=.010, but no differences for fluency factor scores, *p*s > .423). Thus, we may have lost some individuals who were at the greatest risk for later memory impairment, but that would suggest that our findings regarding predictive ability are actually conservative. Strengths of the study are that this is a national, rather than a local, community-based non-clinical sample of middle-aged men with health, education, and lifestyle characteristics that are representative of American men in their age range. The young age of the wave 1 assessment is also a strength with respect to early identification of risk.

## Conclusions

This study demonstrated that episodic memory and semantic fluency independently predicted progression to amnestic MCI in cognitively normal, middle-aged men who were only in their 50s at baseline. Episodic memory also predicted progression to later nonamnestic MCI, although not as strongly as amnestic MCI. These findings demonstrate the usefulness of examining neuropsychological variability among cognitively normal individuals for early identification of risk for AD, highlighting the fact that cognitively normal adults should not be treated as a homogeneous group. Efforts to improve the treatment of AD are beginning to focus on early identification, in part because developing effective treatments are seen as most likely to require intervention during the earliest stages of the disease^13,49^, but also because diagnosing individuals in the stage of MCI could improve patient quality of life and is projected to substantially reduce the financial impact of the disease (e.g., due to non-therapeutic interventions such as financial planning, management of other medical conditions, household safety, etc.)^50^. In the meantime, the results suggest that it will be important to take advantage of the predictive ability of episodic memory and verbal fluency tests in studies designed to identify individuals at greatest risk for MCI (and later dementia). Ultimately, the combination of these cognitive abilities and biomarkers may further improve prediction of MCI and AD.

## Appendix 1 Author Contributions

**Appendix 1: Author Contributions**
| **Name** | **Location** | **Role** | **Contribution** |
| --- | --- | --- | --- |
| Daniel E. Gustavson, Ph.D. | University of California, San Diego | Author | Designed the study; analyzed the data; drafted the manuscript |
| Jeremy A. Elman, Ph.D. | University of California, San Diego | Author | Assisted with data analyses and interpretation; revised the manuscript |
| Matthew S. Panizzon, Ph.D. | University of California, San Diego | Author | Major role in acquisition and interpretation of data; revised the manuscript |
| Carol E. Franz, Ph.D. | University of California, San Diego | Author | Major role in study design, acquisition and interpretation of data; revised the manuscript |
| Jordan Zuber, M.A. | University of California, San Diego | Author | Data interpretation; drafting and revising the manuscript |
| Mark Sanderson-Cimino, M.A. | University of California, San Diego | Author | Data interpretation; revising the manuscript |
| Chandra A. Reynolds, Ph.D. | University of California, Riverside | Author | Data interpretation; revising the manuscript |
| Kristen C. Jacobson, Ph.D. | University of Chicago | Author | Data interpretation; revising the manuscript |
| Hong Xian, Ph.D. | St. Louis University | Author | Data interpretation; revising the manuscript |
| Amy J. Jak, Ph.D. | University of California, San Diego | Author | Data interpretation; revising the manuscript |
| Rosemary Toomey, Ph.D. | Boston University | Author | Major role in acquisition and interpretation of data; revised the manuscript |
| Michael J. Lyons, Ph.D. | Boston University | Author | Major role in study design, acquisition, and interpretation of data; revised the manuscript |
| William S. Kremen, Ph.D. | University of California, San Diego | Author | Major role in study design, acquisition, and interpretation of data; revised the manuscript |

