## Supplemental Information for "Baseline Semantic Fluency Is Associated with Six-Year Progression to Mild Cognitive Impairment in Middle-Aged Men"

*Table S1*

*Demographic Characteristics of the Full Sample*

| Demographic Variable | N | M | SD | Range |
|---|---|---|---|---|
| All Subjects | | | | |
| Lifetime Education | 1483 | 13.8 | 2.11 | 5, 30 |
| Ethnicity (% white non-Hispanic) | 1484 | 88.1 | - | - |
| ApoE status (% e4 positive) | 1444 | 29.4 | - | - |
| Mean age 56 | | | | |
| Age | 1290 | 55.9 | 2.44 | 51.08, 60.67 |
| Depression Symptoms | 1283 | 8.4 | 8.22 | 0, 52 |
| Diabetes (% yes) | 1237 | 11.2 | - | - |
| Hypertension (% yes) | 1237 | 59.8 | - | - |
| Mean age 62 | | | | |
| Age | 1207 | 61.7 | 2.45 | 56.00, 66.92 |
| Age Interval (wave 2 - wave 1) | 1013 | 5.7 | 0.69 | 4.25, 9.42 |

*Note:* Lifetime education was the number of years of school completed. Depression symptoms were measured with the Center for Epidemiologic Studies – Depression Scale[39], with scores above 15 indicating risk for clinical depression. Mean age 56 comprises 1237 individuals ages 51-60 who were tested at study wave 1 plus 53 individuals in this same age range who entered the study and were tested for the first time at wave 2. Mean age 62 comprises 1013 returnees ages 56-66 who were tested a second time at study wave 2 plus 191 attrition replacement subjects in this same age range who entered the study and were tested for the first time at wave 2. The attrition replacement subgroup was recruited specifically to be age-matched to the returnees in order to calculate practice effects for subjects taking the tests a second time. Ns vary in all cases due to missing data.

*Table S2*

*Logistic Regression for Mild Cognitive Impairment (MCI) Predicted by Fluency and Memory Factor Scores with No Covariates*

| Dependent Variable | A) No MCI (N=762) vs. Amnestic MCI (N=42) | | B) No MCI (N=762) vs. Non-Amnestic MCI (N=38) | | C) No MCI (N=762) vs. Any MCI (N=80) | |
|---|---|---|---|---|---|---|
| | OR | 95% CI | OR | 95% CI | OR | 95% CI |
| *Cognitive Factor Scores* | | | | | | |
| General Fluency | 1.27 | [0.80, 2.18] | 1.33 | [0.87, 2.15] | 1.27 | [0.94, 1.77] |
| Semantic-Specific | **1.86** | **[1.15, 3.38]** | 0.97 | [0.63, 1.48] | 1.29 | [0.95, 1.77] |
| Episodic Memory | **4.14** | **[2.38, 9.10]** | **1.95** | **[1.26, 3.32]** | **2.63** | **[1.88, 3.97]** |

*Note:* These models are identical to those displayed in Table 2 of the main text except the covariates are excluded (models still include a random effect to control for the nesting of twins within families). Significant odds ratios (ORs) are displayed in bold ($p < .05$). Factor scores were scored and standardized so the odds ratio indicate the increase in odds of converting to amnestic MCI (A) or non-amnestic MCI (B) at -1 *SD* for that variable. Individuals with non-amnestic MCI were excluded from analyses of amnestic MCI (A) and vice-versa for B. The final column collapses the amnestic and nonamnestic MCI groups into a single “any MCI” group. CI = Confidence interval.

*Table S3*

*Descriptive Statistics for Measures of Verbal Fluency and Episodic Memory in the Full Sample*

| Task | *N* | *M* | *SD* | Range | Skewness | Kurtosis |
|---|---|---|---|---|---|---|
| Mean age 56 | | | | | | |
| *Episodic Memory* | | | | | | |
| Logical Memory | 1279 | 20.01 | 6.63 | 0 - 41 | -0.10 | -0.13 |
| Visual Reproductions | 1283 | 54.75 | 19.51 | 0 - 100 | -0.15 | -0.44 |
| CVLT | 1270 | 9.07 | 2.89 | 0 - 16 | -0.01 | -0.30 |
| *Verbal Fluency* | | | | | | |
| Letter F | 1277 | 12.28 | 4.09 | 1 - 29 | 0.29 | -0.02 |
| Letter A | 1277 | 11.15 | 3.90 | 1 - 29 | 0.41 | 0.34 |
| Letter S | 1277 | 13.48 | 4.32 | 1 - 31 | 0.25 | 0.08 |
| Animals | 1275 | 19.20 | 4.43 | 6 - 39 | 0.26 | 0.26 |
| Boys' Names | 1276 | 19.09 | 4.48 | 6 - 40 | 0.34 | 0.58 |
| Fruits / Furniture | 1277 | 12.75 | 2.55 | 4 - 22 | -0.01 | 0.30 |
| Mean age 62 | | | | | | |
| *Episodic Memory* | | | | | | |
| Logical Memory | 1201 | 17.59 | 6.81 | 0 - 37.36 | -0.09 | -0.35 |
| Visual Reproductions | 1201 | 51.00 | 18.97 | 0 - 95.47 | -0.18 | -0.40 |
| CVLT | 1203 | 8.79 | 2.98 | 0 - 16 | -0.10 | -0.23 |
| *Verbal Fluency* | | | | | | |
| Letter F | 1189 | 11.68 | 4.04 | 2.71 - 21.71 | 0.32 | 0.14 |
| Letter A | 1189 | 10.39 | 3.87 | 1.44 - 26.00 | 0.31 | -0.06 |
| Letter S | 1189 | 12.7 | 4.31 | 0.00 - 28.83 | 0.24 | -0.08 |
| Animals | 1189 | 19.11 | 4.51 | 5.08 - 35.08 | 0.17 | 0.11 |
| Boys' Names | 1189 | 18.32 | 4.5 | 4.31 - 37.31 | 0.23 | 0.47 |
| Fruits / Furniture | 1188 | 12.3 | 2.58 | 2.71 - 21.71 | 0.02 | 0.38 |

*Note*: In all analyses involving the full sample, these dependent measures were standardized residual scores after removing the effect of age on each measure, but the unadjusted scores are presented here. Mean age 56 comprises 1237 individuals ages 51-60 who were tested at study wave 1 plus 53 individuals in this same age range who entered the study and were tested for the first time at wave 2. Mean age 62 comprises 1013 returnees ages 56-66 who were tested a second time at study wave 2 plus 191 attrition replacement subjects in this same age range who entered the study and were tested for the first time at wave 2. The attrition replacement subgroup was recruited specifically to be age-matched to the returnees in order to calculate practice effects for subjects taking the tests a second time. Thus, the scores reported here for the Mean age 62 group reflect the adjustments for practice effects for the returnees. Ns vary in all cases due to missing data.

*Table S4*

*Logistic Regression for Mild Cognitive Impairment (MCI) Predicted by Fluency and Memory Factor Scores (Amnestic and Nonamnestic Collapsed Into Any MCI)*

| Dependent Variable | OR | 95% CI |
|---|---|---|
| General Fluency | 1.24 | [0.91, 1.76] |
| Semantic-Specific | 1.24 | [0.91, 1.71] |
| Episodic Memory | **2.66** | **[1.71, 1.88]** |
| *Covariates* | | |
| Age (wave 1) | **1.50** | **[1.09, 2.15]** |
| Age Interval (wave 2 - wave 1) | 1.17 | [0.85, 1.61] |
| Depression (wave 1) | 0.96 | [0.71, 1.29] |
| ApoE e4+ | 0.86 | [0.43, 1.64] |
| Diabetes (wave 1) | 1.34 | [0.54, 3.18] |
| Hypertension (wave 1) | 1.42 | [0.78, 2.77] |
| Years of Education | 0.88 | [0.63, 1.21] |

*Note:* This model is identical to those displayed in Table 2 of the main text except the amnestic and nonamnestic MCI groups were collapsed into a single any MCI group. Significant odds ratios (ORs) are displayed in bold ($p < .05$). Cognitive factor scores were scored and standardized so the odds ratio indicate the increase in odds of converting to amnestic MCI (A) or non-amnestic MCI (B) at -1 *SD* for that variable. Measures of age, depression, and years of education were also standardized, but not reverse scored. Odds ratios for ApoE status, diabetes, and hypertension reflect increase in odds for having an ε4 allele, diabetes, or hypertension, respectively. CI = Confidence interval. N = 762 (no MCI). N = 80 (any MCI).

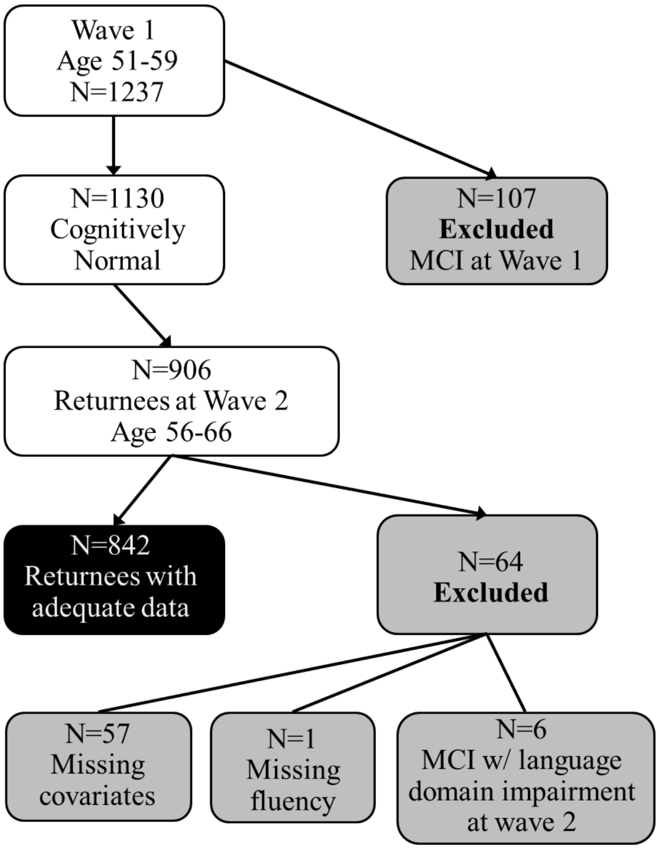

*Figure S1:* The black box indicates final sample of individuals who were cognitively normal at baseline, returned for wave 2, and had all data relevant for this analysis. Gray boxes indicate excluded subjects. Of the 57 subjects missing covariates, 47 were missing the age 20 general cognitive ability measure used in MCI diagnosis, 9 were missing apoE, and 1 was missing depression symptoms.

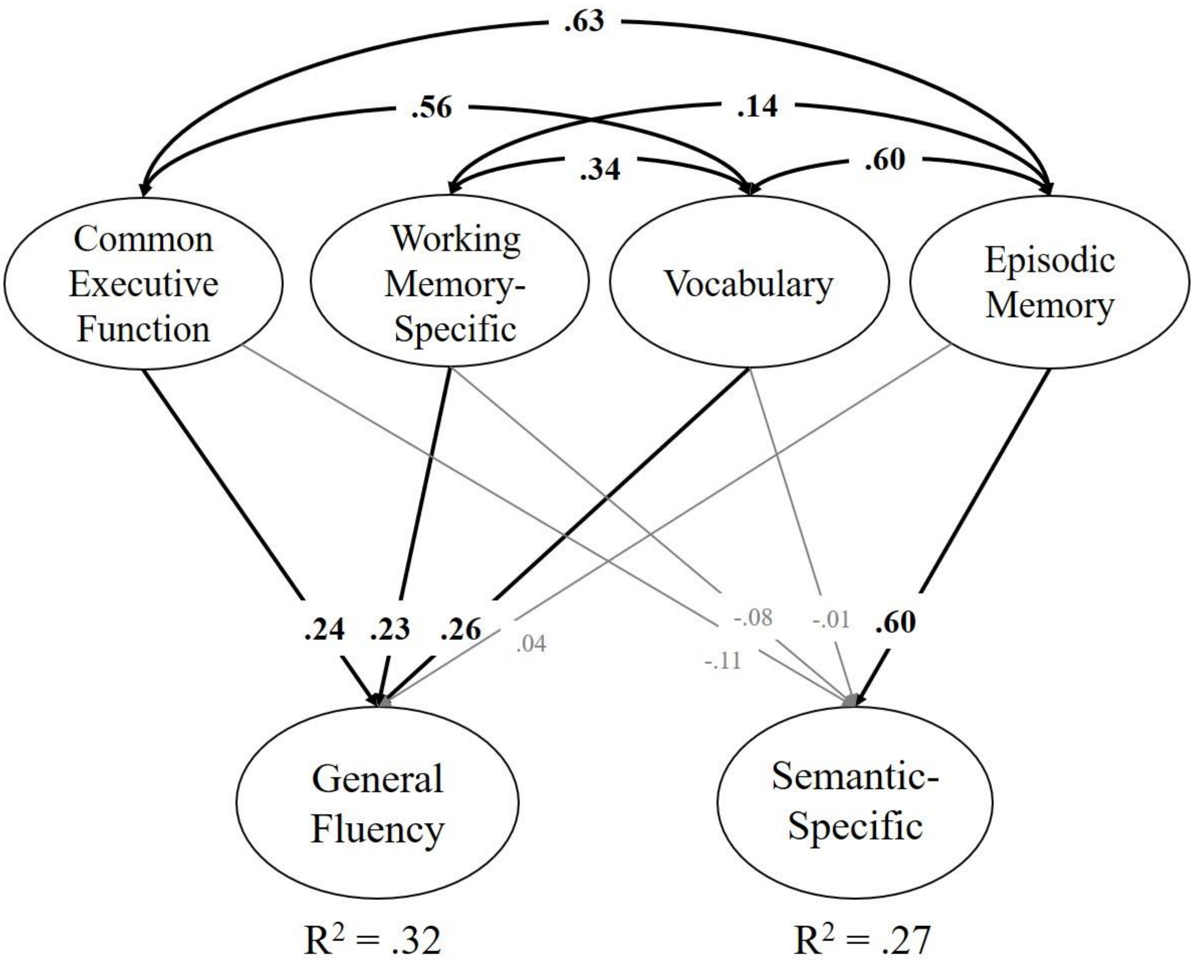

*Figure S2:* Structural equation model where both fluency factors are regressed on latent variables for executive function (Common Executive Function, Working Memory-Specific), vocabulary, and episodic memory. Not pictured are factor loadings on latent factors (which are similar to our previous work and those displayed in Figure 2). Significant paths and correlations are displayed in bold, with black text and lines ($p$ < .05).
